# Single cell RNA-seq denoising using a deep count autoencoder

**DOI:** 10.1101/300681

**Authors:** Gökcen Eraslan, Lukas M. Simon, Maria Mircea, Nikola S. Mueller, Fabian J. Theis

## Abstract

Single-cell RNA sequencing (scRNA-seq) has enabled researchers to study gene expression at a cellular resolution. However, noise due to amplification and dropout may obstruct analyses, so scalable denoising methods for increasingly large but sparse scRNAseq data are needed. We propose a deep count autoencoder network (DCA) to denoise scRNA-seq datasets. DCA takes the count distribution, overdispersion and sparsity of the data into account using a zero-inflated negative binomial noise model, and nonlinear gene-gene or gene-dispersion interactions are captured. Our method scales linearly with the number of cells and can therefore be applied to datasets of millions of cells. We demonstrate that DCA denoising improves a diverse set of typical scRNA-seq data analyses using simulated and real datasets. DCA outperforms existing methods for data imputation in quality and speed, enhancing biological discovery.

## Background

Advances in single-cell transcriptomics have enabled researchers to discover novel celltypes^1,2^, study complex differentiation and developmental trajectories^3–5^ and improve understanding of human disease^1,2,6^.

Despite improvements in measuring technologies, various technical factors including amplification bias, cell cycle effects^7^, library size differences and especially low RNA capture rate lead to substantial noise in scRNA-seq experiments. The low RNA capture rate leads to failure of detection of an expressed gene resulting in a “false” zero count observation, defined as dropout event. Recent droplet-based scRNA-seq technologies, can profile up to millions of cells in a single experiment^8–10^. These technologies are particularly prone to dropout events due to relatively shallow sequencing^11^. Overall, these technical factors introduce substantial noise, which may corrupt the underlying biological signal and obstruct analysis^12^.

In statistics, imputation describes the process of substituting missing data values to improve statistical inference or modeling^13^. However, not all zeros in scRNA-seq data represent missing values. Since not every gene is expected to be expressed in every cell, “true” celltype-specific zeros exist and make the definition of missing values challenging. Therefore, classical imputation methods with defined missing values are not suitable for scRNA-seq data. On the other hand, the concept of denoising, commonly used in image reconstruction^14^, corrects all data entries without first defining a set of missing values.

Current approaches for scRNA-seq specific imputation include scImpute^15^, which defines likely dropout values using a mixture model and subsequently substitutes only the likely dropout values. MAGIC^16^ and SAVER^17^, on the other hand, denoise single-cell gene expression data and generate a denoised output for each gene and cell entry. Despite these differences, all approaches rely on using the correlation structure of single-cell gene expression data to infer “corrected” gene expression values by leveraging similarities between cells and/or genes. With the increasing size of scRNA-seq datasets^11^, methods need to scale to up to millions of cells. However, existing denoising methods are unable to process data sets of this magnitude. Second, linear methods such as scImpute may fail to capture underlying complexity of scRNA-seq.

Therefore, we propose a so-called “deep count autoencoder” (DCA) for denoising scRNA-seq data. An autoencoder is an artificial neural network used for unsupervised learning of the data manifold thereby representing the high dimensional ambient data space in significantly lower dimensions^18^. Ideally, the manifold represents the underlying biological processes and/or cellular states. For example, in a dataset where snapshots of differentiating blood cells exist, the manifold view can show the continuum of differentiation phenotypes. Dimension reduction methods like principal component analysis (PCA), diffusion maps or t-distributed stochastic neighbor embedding (tSNE) are commonly used to visualize the manifold for gene expression data^19,20^. A number of recent studies describe applications of autoencoders in genomics^21–25^. During denoising, the autoencoder learns the manifold and removes the noise by moving corrupted data points onto the manifold^26^ (Fig. 1A). For analogy, PCA can be interpreted as a linear autoencoder with a mean-squared error loss function where the eigenvectors represent the tied encoder and decoder weights. However, due to the complexity and count nature of scRNA-seq data, PCA cannot sufficiently learn the data manifold in many cases^23^. Using an autoencoder out of the box may fail, however, due to the noise model not being adapted to typical scRNA-seq noise. Our DCA approach addresses the challenges underlying scRNA-seq data by 1) enabling non-linear embedding of cells and 2) using scRNA-seq data specific loss function based on negative binomial count distributions (Fig. 1B). We extensively evaluated our approach with competing methods using simulated and real datasets. As expected, we observe increased gene-gene correlation after denoising; while in our examples this enriched for desired regulatory dependencies, this may also lead to overimputation in case of inadequate parameter choices such as too low-dimensional bottleneck layer and hence data manifold. This is a general issue of imputation methods and we propose a hyperparameter search in critical situations. Altogether, we demonstrate that DCA shows high scalability and DCA denoising enhances biological discovery.

**Figure 1.**
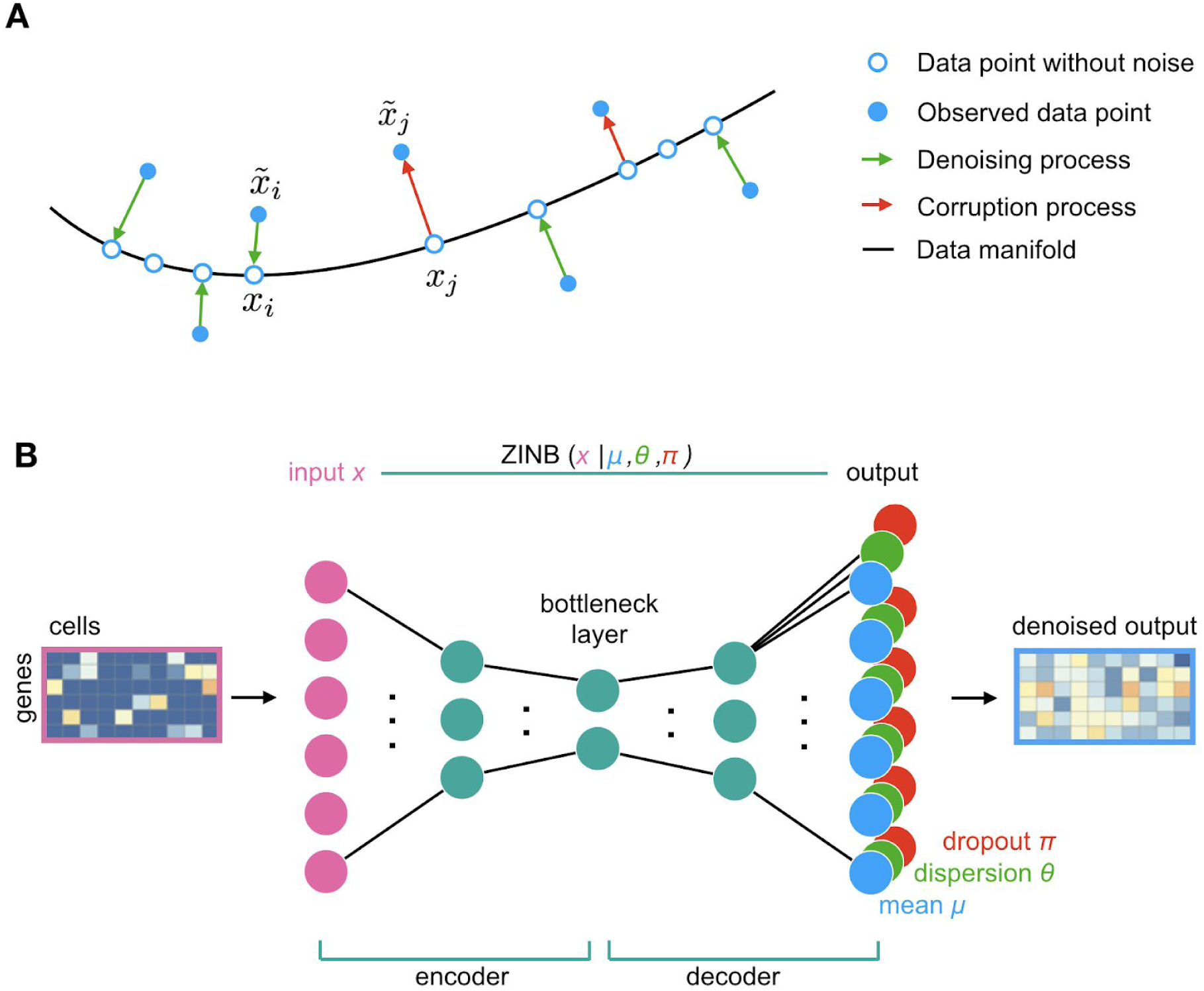
DCA denoises scRNA-seq data by learning the data manifold using an autoencoder framework. Panel A depicts a schematic of the denoising process adapted from Goodfellow et al.^26^. Red arrows illustrate how a corruption process, i.e. measurement noise from dropout events, moves data points ***x̄_j_*** away from the data manifold (black line). The autoencoder is trained to denoise the data by mapping corrupted data points ***x̄_i_*** back onto the data manifold (green arrows). Filled blue dots represent corrupted data points. Empty blue points represent the data points without noise. Panel B shows the autoencoder with a ZINB loss function. Input is the original count matrix (pink rectangle; gene by cells matrix, with dark blue indicating zero counts) and the mean matrix of the negative binomial component represents the denoised output (blue rectangle). Input counts, mean, dispersion and dropout probabilities are denoted as x, μ, θ and π. respectively.

## Results

### Count based loss function is necessary to recover celltypes in simulated scRNA-seq data

Cluster analysis is commonly used in scRNA-seq data to identify celltypes but may be hindered by noise and outliers. To evaluate our method DCA we simulated scRNA-seq datasets including dropout events using Splatter^27^. Both count data with and without dropout are available, which allows quantification of denoising using ground truth. We generated two simulation datasets with 200 genes and 1) two celltypes (2000 cells in total) and 2) six celltypes (2000 cells in total) (see methods for detailed description of simulation parameters). For the two and six celltype simulations 90% and 40% of data values were set to zero, respectively. Dropout simulation probabilities are conditioned on mean gene expression, such that lowly expressed genes have a higher likelihood of dropout compared to highly expressed genes.

Our simulation results show that dropout adds substantial noise, obscuring celltype identities. Expectedly, after denoising using DCA the original celltypes can be recovered (Fig. 2A). To test whether a ZINB loss function is necessary, we compared DCA to a classical autoencoder with mean squared error (MSE) loss function using log transformed count data. The MSE based autoencoder was unable to recover the celltypes, indicating that the specialized ZINB loss function is necessary for scRNA-seq data.

**Figure 2.**
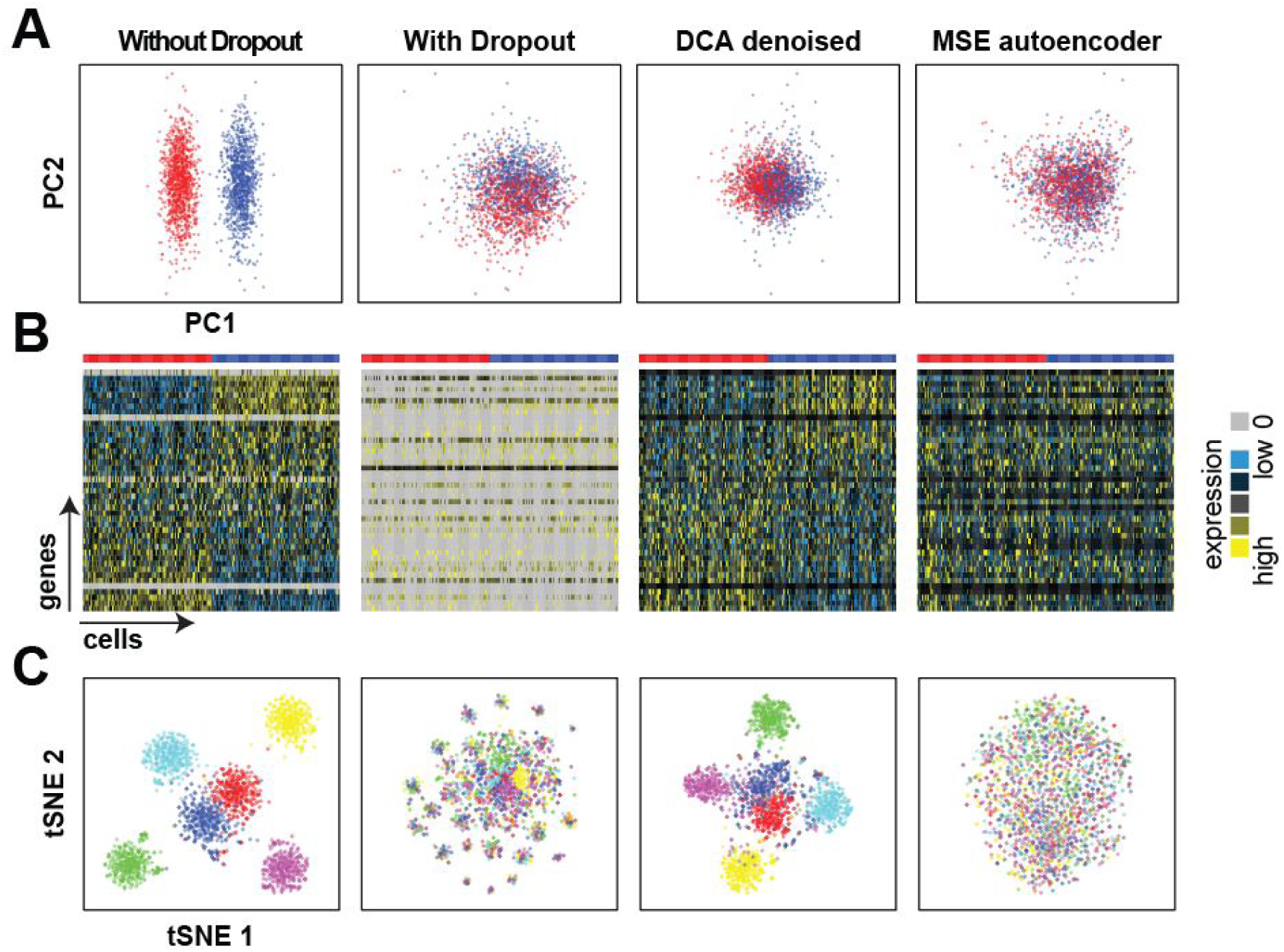
Count based loss function is necessary to identify celltypes in simulated data with high levels of dropout noise. Panel A depicts plots of principal components 1 and 2 derived from simulated data without dropout, with dropout, denoised using DCA and MSE based autoencoder from left to right. Panel B shows heatmaps of the underlying gene expression data. Grey color indicates zero value entries. Panel C illustrates tSNE visualization of simulated scRNA-seq data with six celltypes. Cells are colored by celltype.

Denoising methods bear the risk of introducing spurious correlations, falsely generating correlations between genes. Simulations provide the advantage of defining two sets of genes which 1) show differential expression (DE) between celltypes, i.e. marker genes, and 2) which show no differential expression, i.e. housekeeper genes. Introduction of spurious correlations by denoising could falsely change housekeeper genes into marker genes. To assess whether spurious correlations are introduced by DCA, we performed PCA on the denoised data using the subset of non-DE genes (housekeepers) as input. If denoising introduces spurious gene-gene correlations between marker and housekeeping genes, we expect that PCA on housekeeping genes shows marker gene cluster structure. After DCA denoising, celltype identities were not recovered, indicating that the denoising process did not introduce spurious correlations (Supplementary Fig. 1).

### DCA captures cell population structure in real data

Complex scRNA-seq data sets, such as those generated from a whole tissue, may show large cellular heterogeneity. Celltype marker genes are highly expressed in a celltype-specific manner, leading to “true” celltype-specific zero counts. These are biologically meaningful and need to be distinguished from technical zeros, such as dropout. Therefore, denoising methods must be able to capture the cell population structure and use cell population specific parameters for the denoising process. To test whether DCA was able to capture cell population structure in real data we denoised scRNA-seq data of 68,579 peripheral blood mononuclear cells^10^ and 1,000 highly variable genes (92% zeros) (Fig. 3A). For this analysis only, we restricted the autoencoder bottleneck layer to two neurons and visualized the activations of these two neurons for each cell in a two-dimensional scatter plot (Fig. 3B). When overlaying the original celltype information^10^, celltype-specific clustering was observed. Furthermore, known celltype marker genes showed cluster-specific expression in the two-dimensional bottle neck visualization (Fig. 3 C-F), demonstrating that DCA captures cell population structure in real data. These results prove that DCA can derive cell population specific denoising parameters.

**Figure 3.**
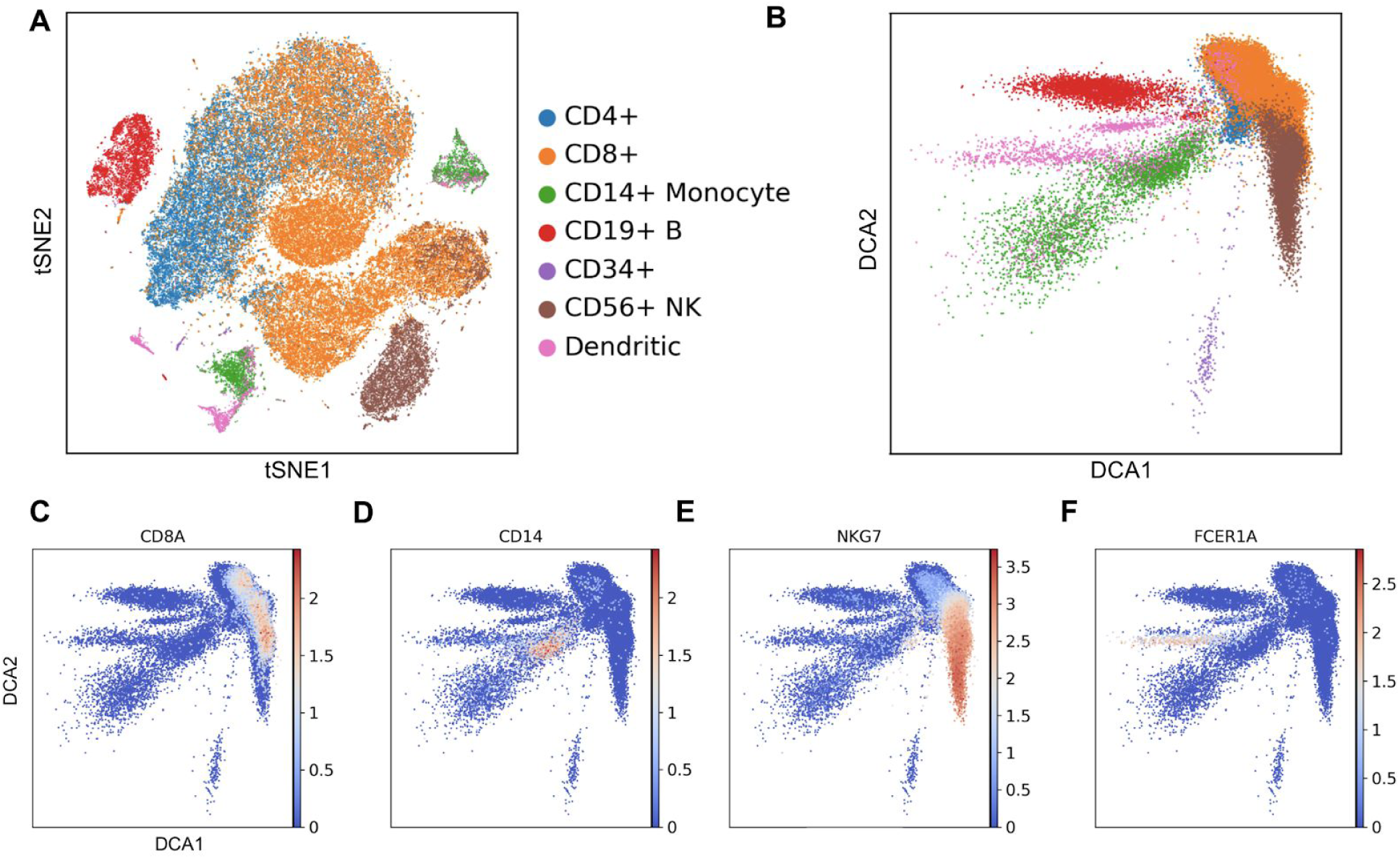
DCA captures population structure in 68,579 peripheral blood mononuclear cells. Panel A shows the tSNE visualization reproduced from Zheng et al.^10^. Panel B illustrates the activations from the two-dimensional bottleneck layer of the DCA. Colors represent celltype assignment from Zheng et al.^10^ where CD4+ and CD8+ cells are combined into coarse groups. Silhouette coefficients are −0.01 and 0.07 for tSNE and DCA visualizations. Panels C-F show two-dimensional bottleneck layer colored by the log-transformed expression of cell type marker genes CD8A (CD8+ T cells), CD14 (CD14+ Monocytes), NKG7 (CD56+ natural killer cells) and FCER1A (dendritic cells), respectively.

### Denoising recovers time-course expression patterns after in silico addition of single-cell specific noise

Next, we evaluated DCA by performing systematic comparison with MAGIC^16^, SAVER^17^ and scImpute^15^ (Supplementary Table 1). We adapted the evaluation approach from van Dijk et al.^16^ and analyzed real bulk transcriptomics data from a developmental C. elegans time course experiment^28^ after simulating single-cell specific noise. Bulk contains less noise than single-cell transcriptomics data^29^ and can thus aid the evaluation of single-cell denoising methods by providing a good ground truth model. Gene expression was measured from 206 developmentally synchronized young adults over a twelve-hour period (Fig. 4A). Single-cell specific noise was added *in silico* by gene-wise subtracting values drawn from the exponential distribution such that 80% of values were zeros^16^ (Fig. 4B). DCA denoising recovered original time course gene expression pattern while removing single-cell specific noise (Fig. 4C). To systematically evaluate the four methods, we tested which method would best recover the top 500 genes most strongly associated with development in the original data without noise. DCA demonstrated strongest recovery of these genes, outperforming the other methods (Fig. 4D). Gene-level expression without, with noise and after DCA denoising for key developmental genes *tbx-36* and *his-8* is depicted in panels E, F, G, respectively. Expression data derived from denoising using MAGIC, SAVER and scImpute for these two genes is displayed in Supplementary Fig. 2. *tbx*-*36* and *his*-*8* represent transcription factor and histone gene classes, respectively, which are known to show opposing expression patterns during C.elegans development^30^.

**Figure 4.**
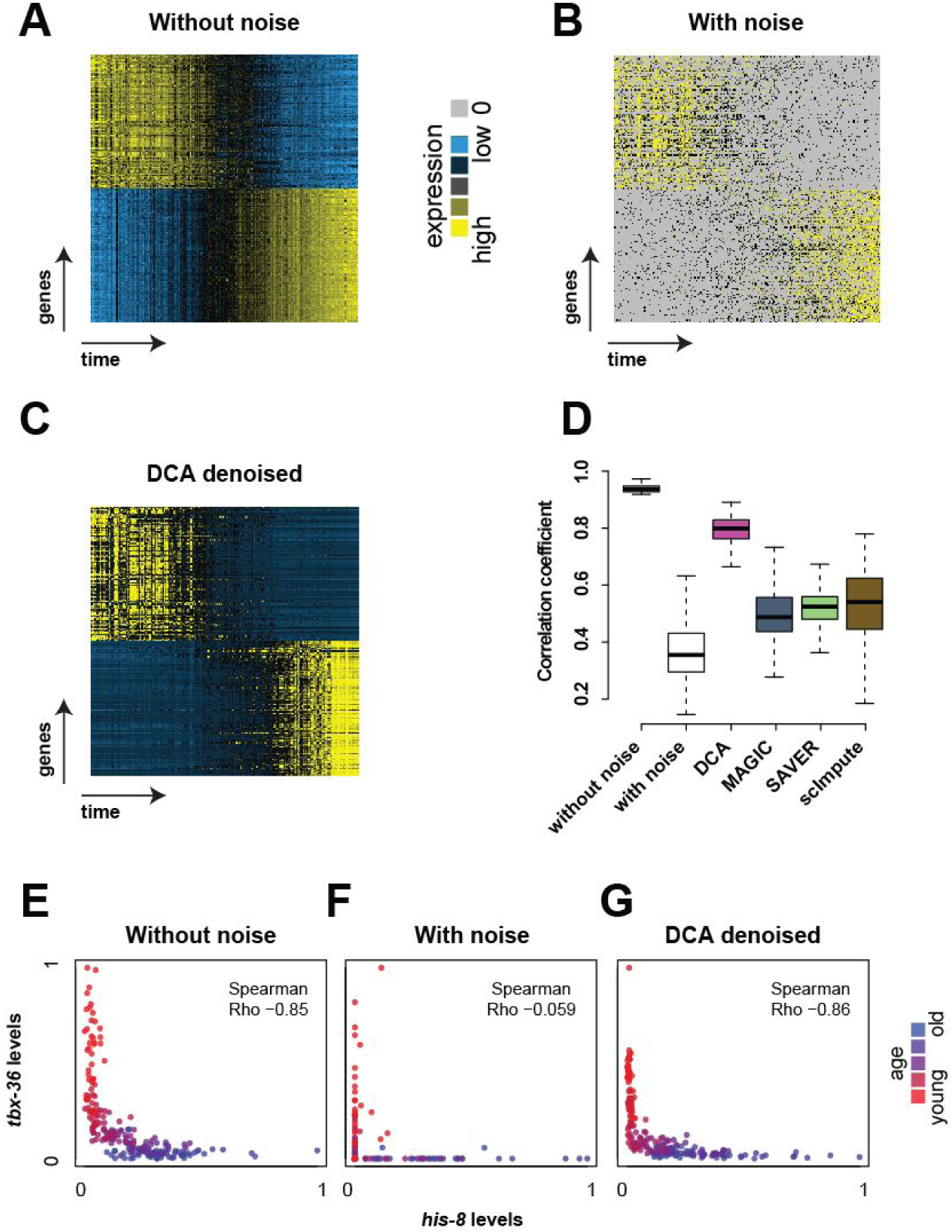
DCA recovers gene expression trajectories in C.elegans time course experiments with simulated dropout. Heatmaps show top 100 genes with positive and negative association with time course using expression data without noise (Panel A), with noise (Panel B) and after DCA denoising (Panel C). Yellow and blue colors represent relative high and low expression levels, respectively. Zero values are colored grey. Distribution of Pearson correlation coefficients across the 500 most highly correlated genes before noise addition for the various expression matrices are depicted in Panel D. The box represents the interquartile range, the horizontal line in the box is the median, and the whiskers represent 1.5 times the interquartile range. Panels E, F and G illustrate gene expression trajectory for exemplary anti-correlated gene pair *tbx*-*36* and *his*-*8* over time for data without, with noise and after denoising using DCA.

### Denoising increases correspondence between bulk and single-cell differential expression

Motivated by the scRNA-seq denoising evaluation metrics proposed by Li et al.^15^, we compared differential expression analysis results between bulk and scRNA-seq data from the same experiment. Chu et al.^31^ generated bulk and scRNA-seq data from H1 human embryonic stem cells (H1) differentiated into definitive endoderm cells (DEC). We performed differential expression analysis comparing H1 to DEC of the bulk and scRNA-seq data independently using DESeq2, which models gene expression based on the negative binomial distribution without zero inflation^32^. DCA slightly increased the Spearman correlation coefficient of the estimated fold changes between bulk and single-cell data from 0.68 to 0.76 (Fig. 5A & B). However, it is important to note that 24 outlier genes (Fig. 5A, red dots), showing high discrepancy between bulk and single-cell derived fold changes, are corrected in the denoised data. *SOX17* is a key transcription factor in the development of the endoderm^33^ and shows high expression in DEC compared to H1 in the bulk data (Fig. C). After DCA denoising, the median expression level of *SOX17* in DEC is shifted higher, more closely reflecting the observation in the bulk data (Fig. 5D-E).

**Figure 5.**
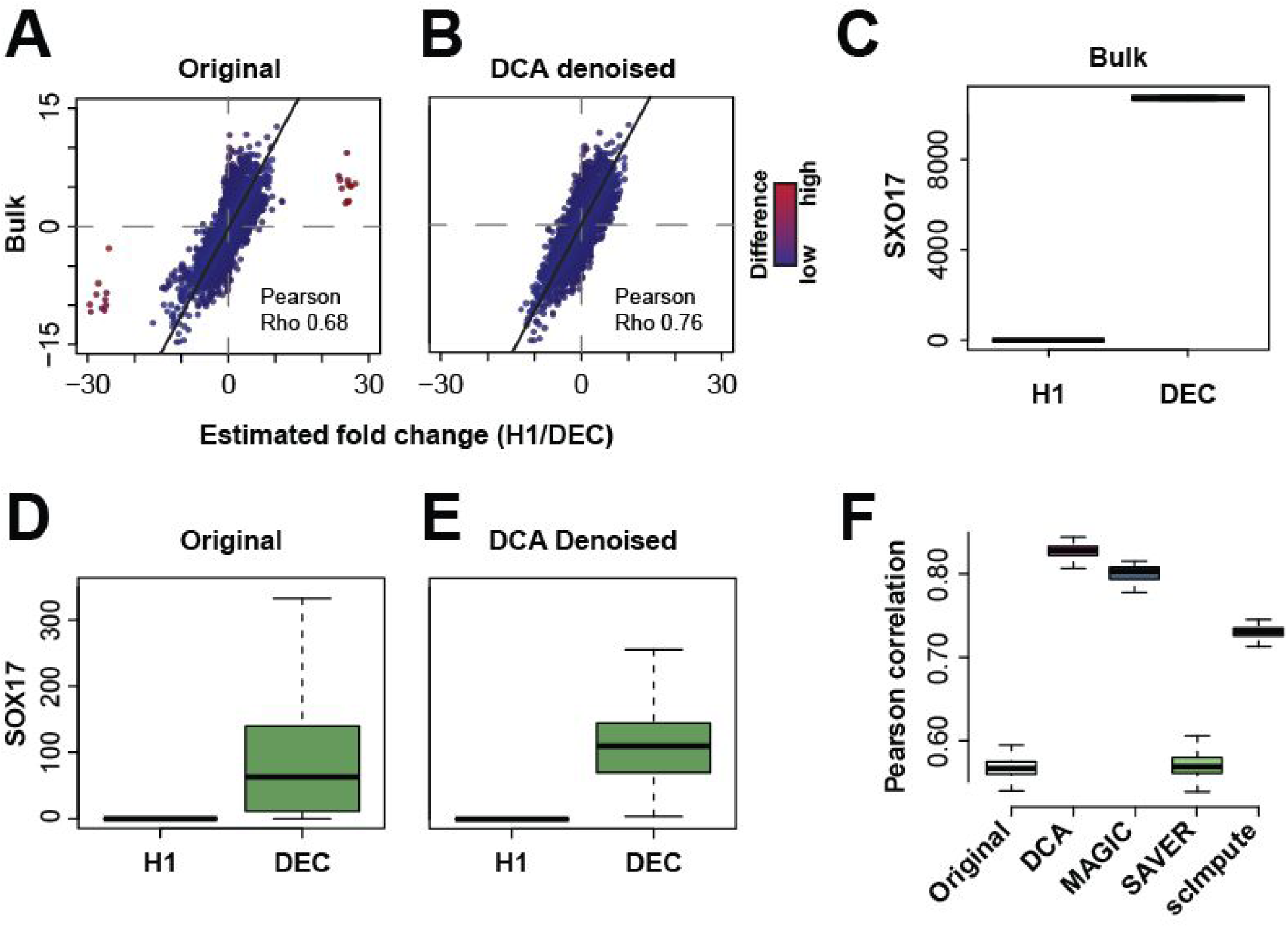
DCA increases correspondence between single cell and bulk differential expression analysis. Scatterplots depict the estimated fold changes for each gene derived from differential expression analysis using bulk and original scRNA-seq count matrix (A), DCA denoised count matrix (B). Grey horizontal and vertical lines indicate zero fold change. Black line indicates identify line. Points are colored by the absolute difference between fold changes from bulk and single-cell data with red colors indicating relative high differences. Panels C, D and E depict differential expression of an exemplary gene *SOX17* between H1 and DEC for the bulk, original and DCA denoised data, respectively. Panel F illustrates boxplots of the distribution of Pearson correlation coefficients from bootstrapping differential expression analysis using 20 randomly selected cells from the H1 and DEC populations for all denoising methods.

Next, we systematically compared the four denoising methods for robustness using a bootstrapping approach. 20 random cells were sampled from H1 and DEC populations one hundred times and differential expression analysis using DESeq2 performed. When comparing the estimated fold changes across all bootstrap iterations, DCA showed highest correspondence with bulk fold changes (Fig. 5F).

### Denoising increases protein and RNA co-expression

CITE-seq enables simultaneous measurement of protein and RNA levels at cellular resolution. Per-cell protein levels are higher than mRNA levels for the corresponding genes and therefore less prone to dropout events^34^. Therefore, by using cell surface marker protein expressions as ‘ground truth’, denoising of mRNA levels can be evaluated. Stoeckius et al. used this CITE-seq method to profile cord blood mononuclear cells and identified major immunological celltypes (Fig. 6A). The original RNA count data was denoised using all four methods and evaluated. Fig. 6B shows tSNE visualization of the data colored by the expression levels of proteins CD3, CD11c, CD56 and corresponding RNAs *CD3E*, *ITGAX*, *NCAM1* by column, respectively. The rows correspond to the protein expression levels, RNA expression levels derived from the original and DCA denoised data. Visualizations for additional protein-mRNA pairs and other methods can be found in Supplementary Fig. 3 and 4, respectively. For example, the CD3 protein is expressed in 99.9% of T cells. The corresponding RNA *CD3E*, however, is only detected in 80% of T cells in the original count data. After denoising using DCA, *CD3E* is expressed in 99.9% of all T cells (Fig. 6C). Some slight discrepancies between the protein and denoised expression can be observed. For example, in the denoised data *ITGAX* shows expression in the natural killer cells (NK) cell cluster while the corresponding CD11c protein levels are very low. Data from the Immunological Genome project (immgen.com), confirmed expression of *ITGAX* in NK cells (data not shown), indicating that the denoised data for this gene reflects better agreement with known biology which may be obscured in the CITE-seq protein data due to some unknown technical reasons. To statistically evaluate the denoising methods we performed co-expression analysis using Spearman correlation for all eight available protein-mRNA pairs across all cells. DCA showed highest median correlation coefficient, indicating that denoising increases protein and RNA co-expression (Fig. 6D).

**Figure 6.**
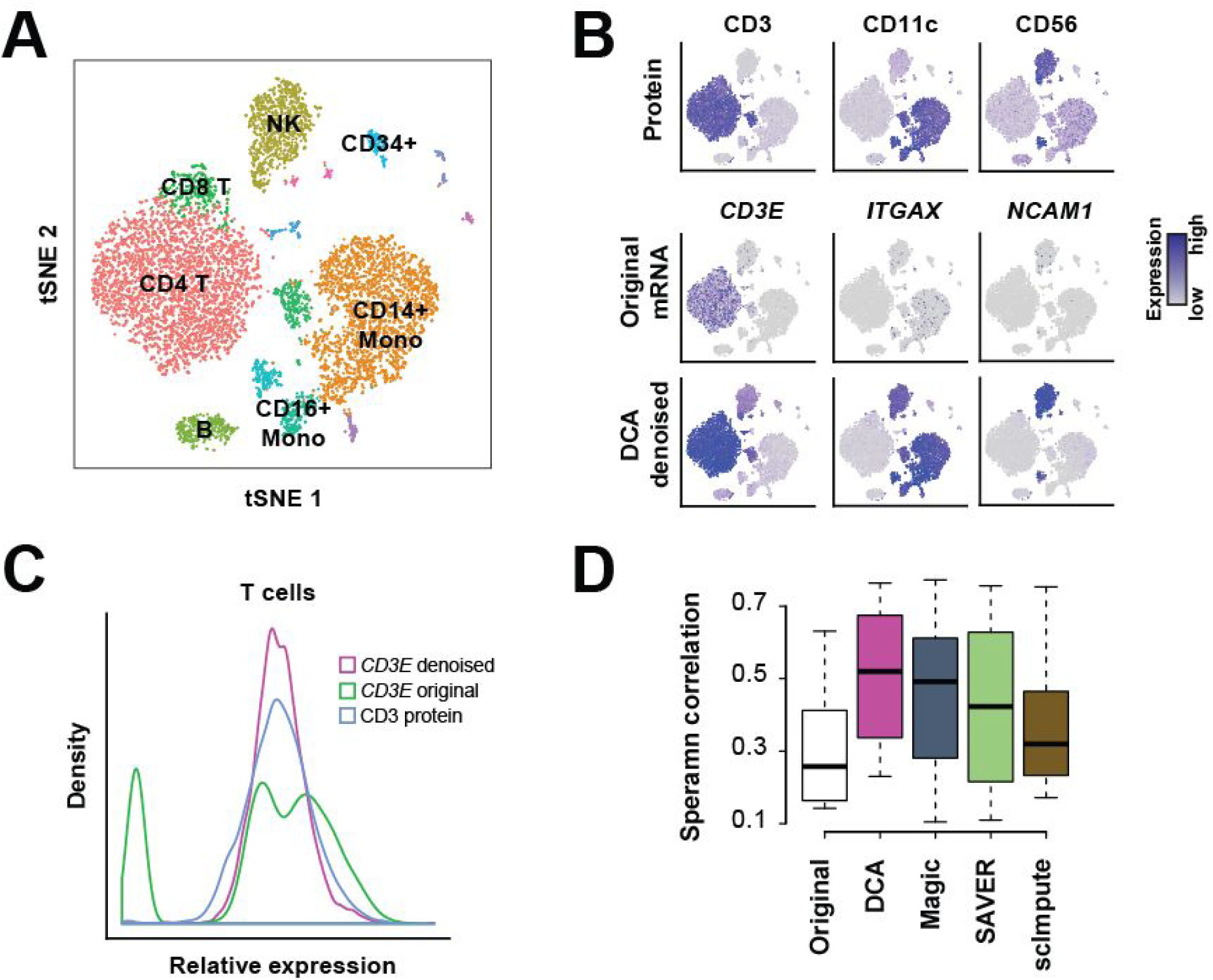
DCA increases protein and RNA co-expression. Panel A depicts tSNE visualization of transcriptomic profiles of cord blood mononuclear cells from Stoeckius et al. Cells are colored by major immunological celltypes. Panel B contains tSNE visualizations colored by protein expression (first row), RNA expression derived from original (second row) and DCA denoised data (third row). Columns correspond to CD3 (first column), CD11c (second column), CD56 (third column) proteins and corresponding RNAs *CD3E*, *ITGAX* and *NCAM1*. Panel C shows the distribution of expression values for CD3 protein (blue), original (green) and DCA denoised (pink) *CD3E* RNA in T cells. Spearman correlation coefficients for the eight protein-RNA pairs across all cells for the original and denoised data are plotted panel D.

### DCA runtime scales linearly with the number of cells

As the number of cells profiled in a single experiment is increasing, it is essential that scRNA-seq methods show good scalability. To assess the scalability of the four methods, we analyzed the currently largest scRNA-seq data set, consisting of 1.3 million mouse brain cells, from 10X Genomics. The 1.3 million cell data matrix was downsampled to 100, 1,000, 2,000, 5,000, 10,000 and 100,000 cells and 1000 highly variable genes. Each subsampled matrix was denoised and the runtime measured (Fig. 7). The runtime of DCA scaled linearly with the number of cells. While it took DCA minutes to denoise 100,000 cells, the other methods took hours. Therefore, DCA possesses a considerable speed advantage over the competing methods.

**Figure 7.**
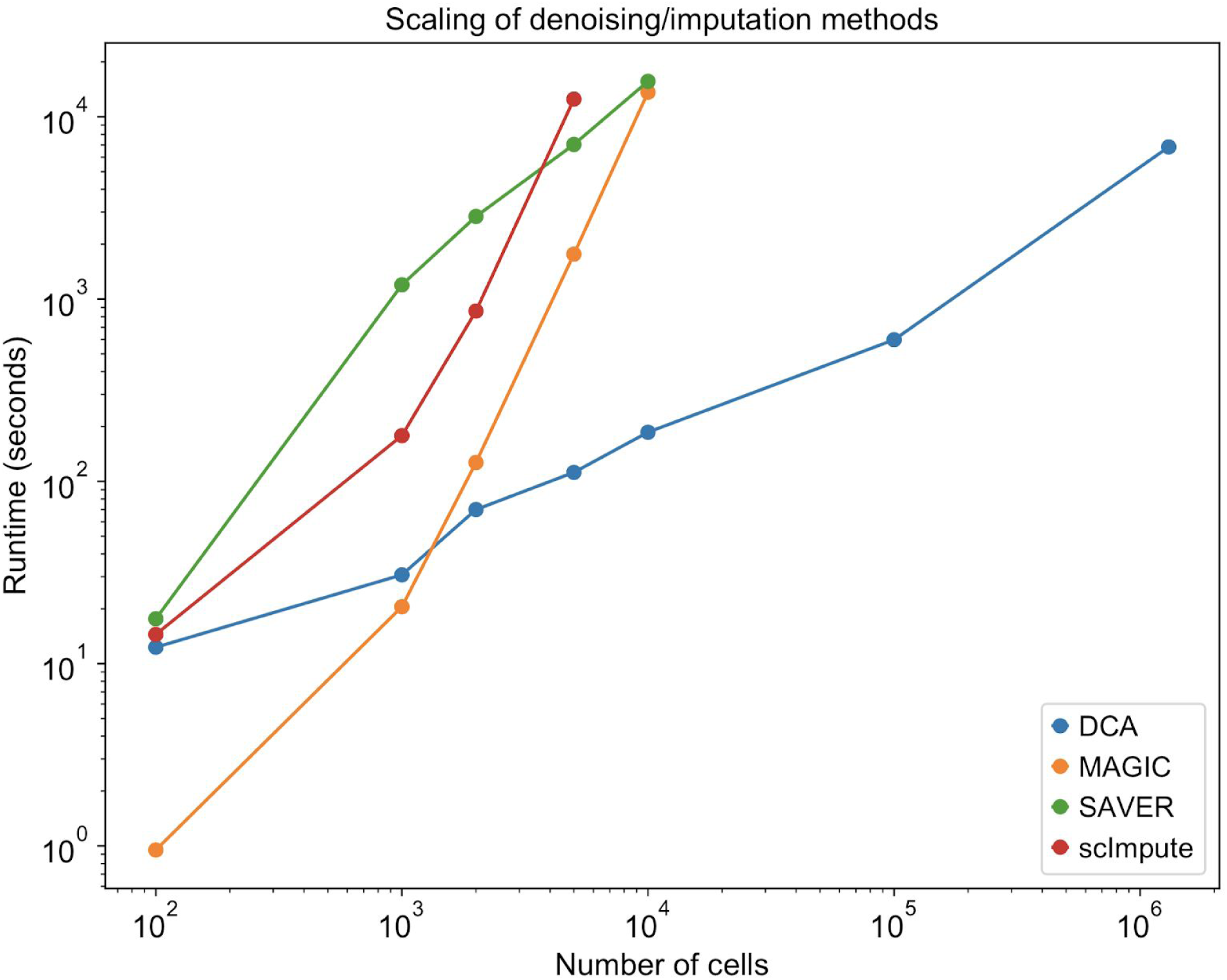
DCA scales linearly with the number of cells. Plot shows the runtimes for denoising of various matrices with different numbers of cells down-sampled from 1.3 million mouse brain cells^35^. Colors indicate different methods.

### Denoising by DCA enables discovery of subtle cellular phenotypes

After having evaluated DCA against competing methods, we tested if DCA denoising could enhance biological discovery which is impossible or more challenging to obtain without denoising. Stoeckius et al.^34^ highlight the potential for integrated and multimodal analyses to enhance discovery of cellular phenotypes, particularly when differentiating between cell populations with subtle transcriptomic differences. The authors observed an opposing gradient of CD56 and CD16 protein levels within the transcriptomically derived NK cell cluster (Fig. 8 A & B). Indeed, unsupervised clustering using Gaussian mixture model on the CD16 and CD56 protein expression levels revealed two sub-populations of cells (Fig. 8C). The corresponding RNAs *NCAM1* and *FCGR3A*, however, contained high levels of dropout obscuring the sub-population structure (Fig. 8D). After denoising, the two sub-populations of NK cells become evident solely based on DCA denoised *NCAM1* and *FCGR3A* RNA expression levels. Therefore, DCA denoising enabled the extraction of information which was exclusively contained in the CITE-seq proteins, demonstrating the ability to enable discovery of subtle cellular phenotypes (Fig. 8E).

**Figure 8.**
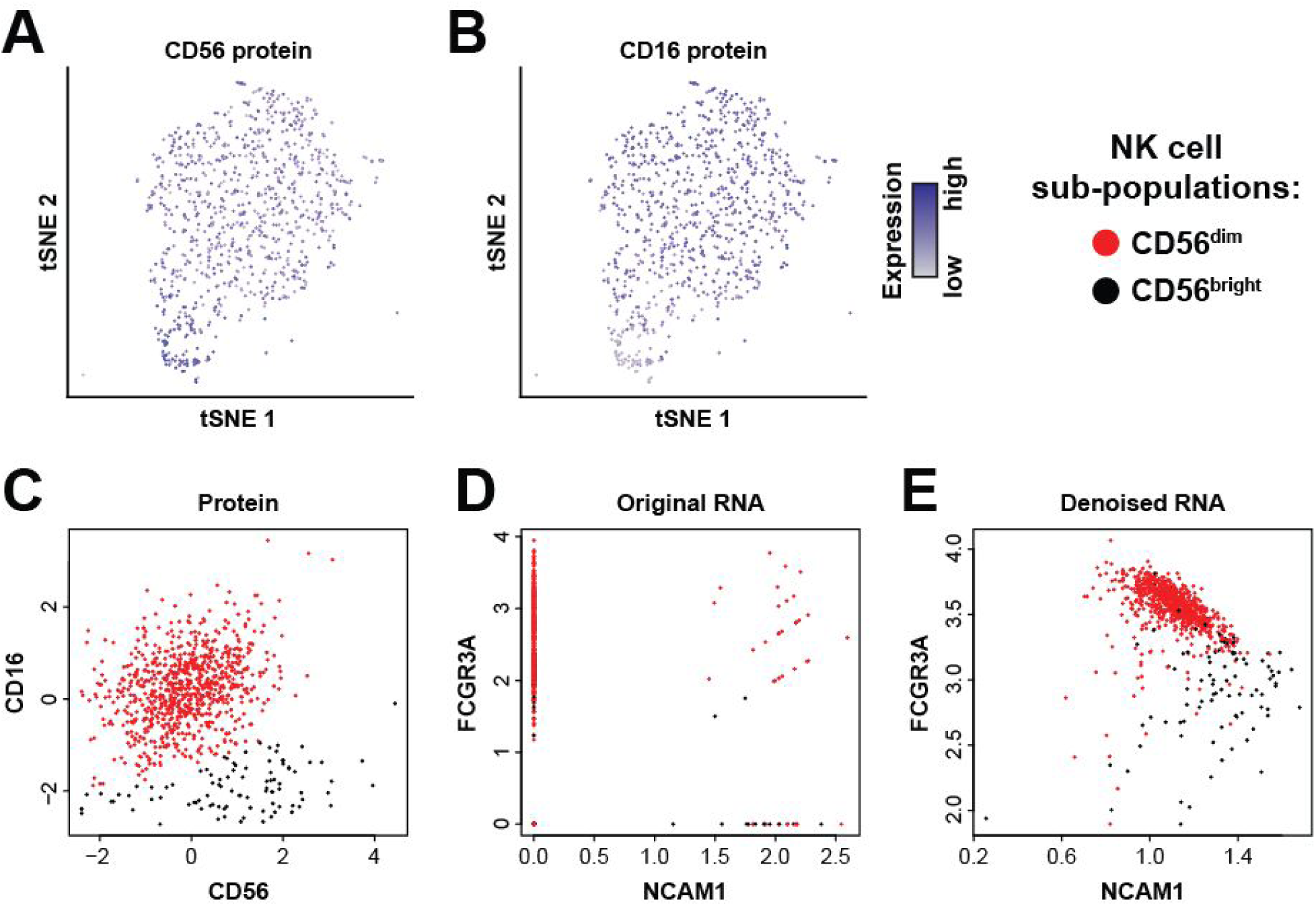
Denoising enhances discovery of cellular phenotypes. tSNE visualization of transcriptomically derived NK cell cluster colored by CD56 (panels A) and CD16 (panel B) protein expression levels. Grey and blue indicate relative low and high expression, respectively. Panel C shows CD56 and CD16 protein expression across NK cells, revealing two distinct sub-populations defined as CD56dim (red) and CD56bright (bright). Panels D and E depict expression of corresponding RNAs *NCAM1* and *FCGR3A* using the original count data and DCA denoised data, respectively. Cells are colored by protein expression derived assignment to CD56bright (black) and CD56dim (red) NK cell sub-populations.

### Denoising by DCA increases correlation structure of key regulatory genes

Next, we tested if denoising enhances discovery of regulatory relationships for well-known transcription factors in blood development. In Paul et al.^36^ the authors describe the transcriptional differentiation landscape of blood development into megakaryocyte–erythroid progenitors (MEP) and granulocyte-macrophage progenitors (GMP) (Fig. 9A-B). After denoising, a set of well-known MEP and GMP regulators^37^ show enhanced regulatory correlations (Fig. 9C-D), for example, the anticorrelation between *Pu.1* and *Gata1* increases (Fig. 9E-F). These two transcription factors are important regulators in blood development and known to inhibit each other^38^. This regulatory relationship is identified in denoised data also in cells with zero expression for either gene in the original data, demonstrating the ability of DCA to extract meaningful information from otherwise non-informative zero count values (Supplementary Fig. 5). Overall these results demonstrate that DCA enhances modeling of gene regulatory correlations, and we expect future network inference methods to use denoising as a first preprocessing step.

**Figure 9.**
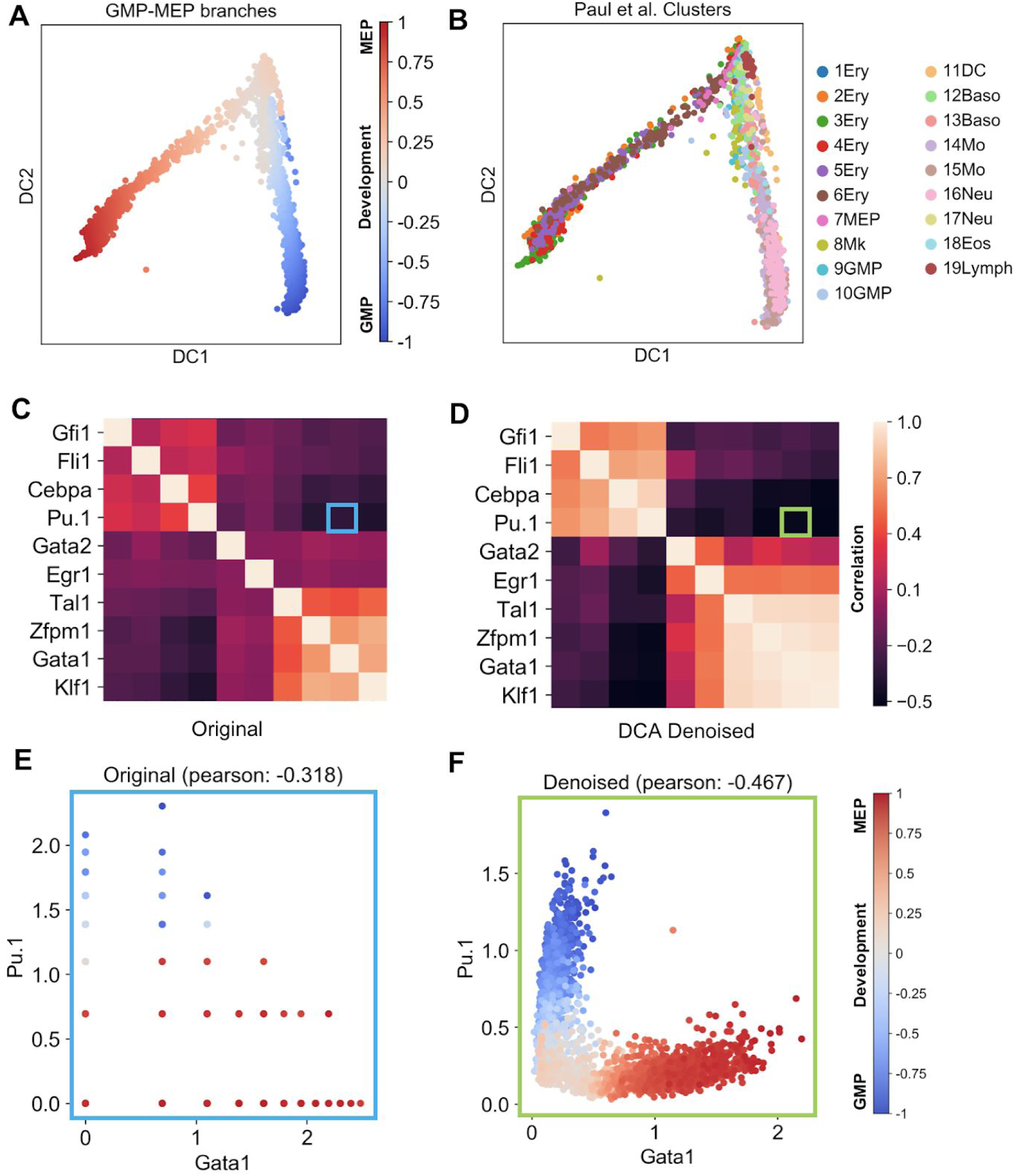
Denoising by DCA increases correlation structure of key regulatory genes. Panels A and B display diffusion maps of blood development into GMP and MEP colored by developmental trajectory and celltype, respectively. Abbreviations Ery, Mk, DC, Baso, Mo, Neu, Eos, Lymph correspond to erythrocytes, megakaryocytes, dendritic cells, basophils, monocytes, neutrophils, eosinophils and lymphoid cells, respectively. Panels C and D display heatmaps of correlation coefficients for well-known blood regulators taken from Krumsiek et al.^37^. Highlighted areas show *Pu. 1* - *Gata1* correlation in the heatmap. Panels E and F show anti-correlated gene expression patterns of *Gata1* and *Pu.1* transcription factors colored by pseudotime, respectively.

## Discussion

One of the fundamental challenges in scRNA-seq analysis is technical variation. Recent research has shown that accounting for technical variation improves downstream analysis^7,39–11^ such as uncovering the cell differentiation structure, identification of highly variable genes, and clustering. Furthermore, some denoising/imputation methods have been implemented in scRNA-seq workbenches such as Granatum^42^, indicating that it is an important, frequently used processing or smoothing step e.g. for visualization.

Here, we introduce a robust and fast autoencoder-based denoising method tailored to scRNA-seq datasets. We demonstrate that denoising scRNA-seq data can remove technical variation improving five possible downstream analyses, namely clustering, time course modeling, differential expression, protein-RNA co-expression and pseudotime analyses.

The evaluation of denoising is difficult because the definition of a ground truth can be challenging for real data. We therefore described a diverse set of evaluation scenarios, which may allow systematic assessment of other denoising techniques in the future. Furthermore, in order to avoid bias in comparisons, we adapted evaluation approaches and used corresponding data from competing methods for evaluation.

Note that in general it may be difficult to determine when denoising improves scRNA-seq data and careful estimation of overimputation may be necessary, for example by hyperparameter optimization or regularization. To alleviate overfitting and overimputation, a general and not yet extensively treated issue of imputation methods, we implemented a number of regularization methods, including dropout, encoder-specific and overall L1 and L2 regularization. This is required especially when training on data sets with limited sample size. DCA also allows users to conduct a hyperparameter search to find the optimal set of parameters for denoising to avoid poor generalization due to overfitting.

The proposed method can be easily integrated into existing workflows; in particular it supports h5ad-formatted HDF5 files (https://github.com/theislab/anndata) and the Python API is compatible with Scanpy^43^ package. Furthermore, we show that DCA is highly scalable to data sets with up to millions of cells.

## Methods

### Architecture and training

Zero-inflated negative binomial (ZINB) is a distribution used for modelling highly sparse and overdispersed count data. ZINB mixture model consists of two components, a point mass at zero which represents excess zeros in datasets and a negative binomial component representing the count process. For the scRNA-seq data, the point mass corresponds to dropout zeros whereas the negative binomial component represents the process that gives rise to the count data.

ZINB is parameterized with mean and dispersion parameters of the negative binomial component (μ and θ) and the mixture coefficient that represents the weight of the point mass (π):

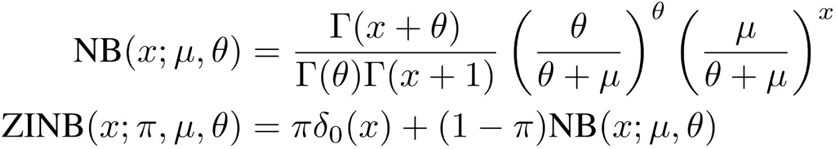

A typical autoencoders compresses high dimensional data into lower dimensions in order to constrain the model and extract features that summarize the data well in the bottleneck layer. In scRNA-seq context, these features are ideally cell type specific. The hidden features are then used by the decoder to estimate the mean parameter of a normal distribution for each feature i.e. each gene in scRNA-seq context. Therefore, there is a single output layer representing the mean. Here, we use the autoencoder framework to estimate three parameters of ZINB distribution conditioned on the input data for each gene. Therefore, unlike traditional autoencoders, our model also estimates dropout (π) and dispersion (θ) parameters in addition to the mean (μ). Each module corresponds to a parameter of the ZINB distribution, given as μ, θ and π. In binary classifiers, the output layer is interpreted as logistic regression using the features extracted from the previous layers. Similarly, the output layer in our approach can be interpreted as ZINB regression where predictors are new representations of cells. The formulation of the architecture is given below:

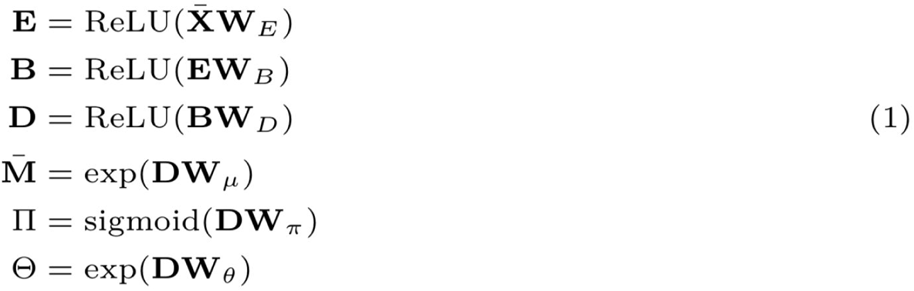

where **E**, **B** and **D** represent encoder, bottleneck and decoder layers, respectively. In this formulation, **X̄** represents library size, log and z-score normalized expression matrix where rows and columns correspond to cells and genes, respectively. Size factors for every cell, S_j_, is calculated as total number of counts per cell divided by the median of total counts per cell. **X̄** is defined as:

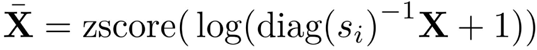

where **X** and *zscore* represent the raw count matrix and z-score normalization.

Output activations are shown here in matrix form as **M̄**, Θ and **Π**. Although the mini-batch stochastic gradient descent is used for optimization, for convenience here we depict the entire matrices of size n × p where n and p represent number of cells and genes, respectively.

Note that in the architecture, the activation function of the mean and dispersion output layers is exponential, since the mean and dispersion parameters are always non-negative. The third output **Π** estimates the dropout probability for every element of the input. The activation function of this layer is sigmoid as **Π** values represent the dropout probabilities. The activation function of the three output layers is an inverse canonical link function of a ZINB regression model in the context of generalized linear models.

The loss function that is likelihood of zero-inflated negative binomial distribution:

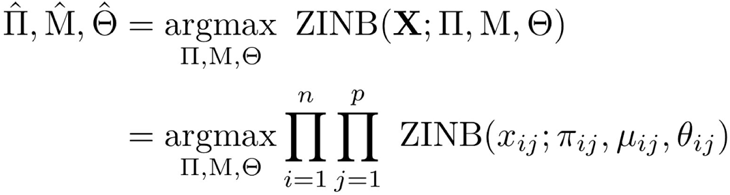

where x_ij_ represents the elements in the raw count matrix **X**. i and j represent cell and gene indices and n and p represent the number of cells and genes. **M** represents the mean matrix multiplied by the size factors that are calculated before the training:

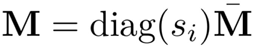

which keeps the hidden representation of cells and the optimization process independent of library size differences.

Furthermore, our model contains a tunable zero-inflation regularization parameter that acts as a prior on the weight of the dropout process. This is achieved by the ridge prior over the dropout probabilities and zero inflation (**Π** parameter):

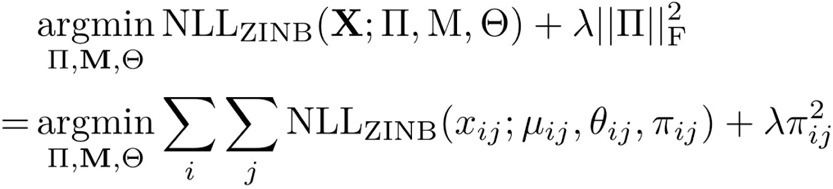

where NLL_ZINB_ function represents the negative log likelihood of ZINB distribution.

To increase flexibility we provide implementations of a negative binomial loss function with (ZINB) and without zero-inflation (NB). Hyperparameter search allows users to find optimal λ value for given dataset. Furthermore, users are also allowed to choose whether the dispersion parameter is conditioned on the input. While n × p dispersion matrix is estimated from the data in the conditional dispersion (default option), the alternative option estimates a scalar dispersion parameter per gene.

### Denoising

The denoised matrix is generated by replacing the original count values with the mean of the negative binomial component (**M̄** matrix in Equation 1) as predicted in the output layer. This matrix represents the denoised and library size normalized expression matrix, the final outcome of the method. Intuitively, our approach can be interpreted as a two-step process. First, the data is summarized by extracting lower dimensional hidden features that are useful for denoising the data as well as identifying and correcting dropout zeros. Then, a ZINB regression is fitted using these new hidden features. However, these two steps are performed simultaneously during the training.

### Implementation

DCA is implemented in Python 3 using Keras^44^ and its TensorFlow^45^ backend. We used RMSProp for optimization with learning rate 0.001. Learning rate is multiplied by 0.1 if validation loss does not improve for 20 epochs. The training stops after no improvement in validation loss for 25 epochs. Gradient values are clipped to 5 and the batch size is set to 32 for all datasets. All hidden layers except for the bottleneck consist of 64 neurons. Bottleneck has 32 neurons. Training on CPU and GPU is supported thanks to Keras and TensorFlow.

Hyperparameter search is implemented using hyperopt^46^ and kopt Python packages (https://github.com/Avsecz/koph. One thousand hyperparameter configurations such as different number of layers, number of neurons, L1/L2 coefficients are trained for 100 epochs using the Tree-structured Parzen Estimator (TPE)^46^ method implemented in hyperopt. Model with lowest validation error is reported.

### Simulated scRNA-seq data

Simulated data sets were generated using the Splatter R package^27^. For the two group simulation the following parameters were used in the *splatSimulate()* R function groupCells = 2, nGenes = 200, dropout.present = TRUE, dropout.shape −1, dropout.mid = 5. For the six group simulation the following parameters were used in the *splatSimulate()* R function groupCells = 6, nGenes = 200, dropout.present = TRUE, dropout.shape −1, dropout.mid = 1.

### 68k peripheral blood mononuclear cell experiment

Single cell gene expression count matrix and celltype labels from Zheng et al. were downloaded from http://www.github.com/10XGenomics/single-cell-3prime-paper. Since CD4+ and CD8+ subtype clusters are highly overlapping, they are combined into coarse groups. tSNE coordinates were obtained by reproducing the code from single-cell-3prime-paper repository. DCA was run using the following parameter: -s 16,2,16.

### MAGIC

MAGIC was downloaded from https://github.com/KrishnaswamvLab/maaic. MAGIC was run using default parameters specified as 20 for the numbers of principal components, 6 for the parameter t for the power of the markov affinity matrix, 30 the number of nearest neighbors, 10 the autotune parameter and 99th percentile to use for scaling.

### sclmpute

sclmpute (version 0.0.5) was downloaded from https://github.com/Vivianstats/sclmpute. For the comparison the following parameters were used kCluster = 2. For the CITE-seq cord blood mononuclear cells experiment and the scalability analysis, kCluster =13 and kCluster = 2 parameters are used, respectively.

### SAVER

SAVER (version 0.3.0) was downloaded from https://github.com/mohuangx/SAVER. SAVER was run using default parameters specified as 300 for the maximum number of genes used in the prediction, 50 for the number of lambda to calculate in cross validation and 5 for the number of folds used in cross validation. For the scalability analysis, SAVER was run using the R package *doParallel* with 24 cores.

### DCA

For the two group simulation data the following parameters were used --*type zinb*-*cond*. For the six group simulation data the following parameters were used --*type zinb*. For the C. elegans development experiment the following parameters were used --*type zinb*. For the Chu et al. definitive endoderm differentiation experiment the following parameters were used --*type nb*. For the CITE-seq cord blood mononuclear cells experiment, the following parameters were used --*type nb*.

### C. elegans development experiment

Bulk microarray gene expression of developing C.elegans embryos was downloaded the supplementary material of Francesconi et al.^28^. This data set contained 206 samples covering a 12-hour time-course. Similar to the evaluation proposed van Dijk et al., expression values were exponentiated to create a count distribution and subsequently single-cell noise was added *in silico* by subtracting gene-specific artificial noise from each gene. Gene-specific artificial noise was generated using the exponential function where the mean was calculated as the gene expression median multiplied by five. Any negative values were set to zero so that on average 80% of the values were zero. Pearson correlation was calculated between the expression level of each gene and time course to identify top 500 development genes.

### Chu et al. definitive endoderm differentiation experiment

Single-cell and bulk RNA-seq data were downloaded from the Gene Expression Omnibus (GEO) accession GSE75748. The gene expression data was restricted to single cells and bulk samples from H1 and DEC using the provided annotation. Differential expression analysis was performed using the R package DESeq2 (version 1.14.1). DESeq2 models gene expression based on a negative binomial distribution without zero-inflation. To retain count structure, denoised data for all methods was rounded prior to analysis. Dispersion was estimated using “mean” for the *fitType* parameter. To assess robustness of the results, bootstrapping analysis was conducted. During each of 100 iterations, 20 cells from the H1 and DEC cells were randomly selected and differential expression analysis performed as described above. Next, concordance was evaluated using Pearson correlation between the estimated fold changes derived from the single-cell bootstrap and bulk data.

### CITE-seq cord blood mononuclear cells experiment

Single cell protein and RNA raw count expression matrices were obtained from the GEO accession GSE100866. The Seurat R package was used to perform the analysis. Following the instructions of the authors^34^ data was subset to 8,005 human cells by removing cells with less than 90% human UMI counts. Next, RNA data was normalized, highly variables genes were identified and expression data was scaled. First 13 principal components were calculated and used for clustering and tSNE visualization. A total of 13 clusters were identified. The *genesCLR* method was used for normalization of the protein data. For denoising, gene expression data was restricted to the top 5000 highly variable genes. Co-expression for eight known marker proteins (CD3, CD19, CD4, CD8, CD56, CD16, CD11c, CD14) and corresponding mRNAs (*CD3E*, *CD19*, *CD4*, *CD8A*, *NCAM1*, *FCGR3A*, *ITGAX*, *CD14*) was assessed using Spearman correlation on the scaled expression data across all 8,005 cells.

### NK subset analysis

Stoeckius et al.^34^ data was subset to 906 NK cells. Next, protein and RNA expression data were scaled. Using CD16 and CD56 protein expression levels, cells were clustered with the *Mclust()* function from the R *mclust* package and two mixture components.

### Blood regulator analysis

Paul et al. blood differentiation data with 2730 cells and 3451 informative genes including the cell type annotations are obtained via *” scanpy.api.datasets.paul15()”*function of Scanpy. After log transform with pseudo count of one, kNN graph is constructed with *“scanpy.api.pp.neighbors”* function. Diffusion map, diffusion pseudotime (DPT) and four diffusion pseudotime groups are computed with *“scanpy.api.tl.dpt(adata*, *n*_*branchings=1)”.* Pseudotime estimates of two DPT groups corresponding to MEP and GMP branches are scaled between [0, 1] and [0, −1] in order to show the branching more distinctly. DCA is run with default parameters and Pearson correlation coefficients between marker genes are calculated with *“numpy.corrcoef”* function.

### Scalability analysis

1.3 million mouse brain cell data were downloaded from https://support.10xaenomics.com/single-cell-aene-expression/datasets/1.3.0/1M_neurons. First, cells and genes with zero expression are removed from the count matrix. Next, top one thousand highly variable genes are selected using “filter_genes_dispersion” function of Scanpy with n_top_genes=1000 argument. The 1.3 million cell data matrix was downsampled to 100, 1,000, 2,000, 5,000, 10,000 and 100,000 cells and these 1000 highly variable genes. Each subsampled matrix was denoised with the four methods and the runtime measured. Scalability analysis was performed on a server with two Intel Xeon E5-2620 2.40GHz CPUs.

### Data Availability

DCA, including usage tutorial, can be downloaded from https://github.com/theislab/dca.

## List of abbreviations

scRNA-seq: single-cell RNA sequencing
tSNE: t-distributed stochastic neighbor embedding
DCA: deep count autoencoder
AE: autoencoder
PCA: principal component analysis
H1: human embryonic stem cells
DEC: definitive endoderm cells
MEP: megakaryocyte-erythroid progenitors
GMP: granulocyte-macrophage progenitors
MSE: mean squared error
ZINB: zero-inflated negative binomial
CITE-seq: Cellular Indexing of Transcriptome and Epitopes by sequencing
NK: natural killer cells
DPT: diffusion pseudotime
ReLU: rectified linear unit.

## Contributions

FJT designed the research. GE, LS and MM carried out the data analysis. FJT, NSM, LS and GE contributed to the manuscript.

## Corresponding author

Correspondence to Fabian J. Theis

